# Life-span development of functional brain networks as assessed with minimum spanning tree analysis of resting state EEG

**DOI:** 10.1101/022038

**Authors:** Dirk J.A. Smit, Eco J.C de Geus, Maria Boersma, Dorret I. Boomsma, Cornelis J. Stam

**Author notes:** Corresponding author: Dr. D.J.A. Smit., Biological Psychology, van der Boechorststraat 1, 1081 BT Amsterdam, The Netherlands.

## Abstract

The brain matures with large quantitative changes in anatomy and function. Graph analysis of EEG has previously revealed increased connectivity between distant brain areas and a decrease in randomness and increased integration in the brain network with concurrent increased modularity. Comparisons of graph parameters across age groups, however, may be confounded with network degree distributions. Here, we analyzed graph parameters from minimum spanning tree (MST) graphs. MST graphs are constructed by selecting only the strongest available connections avoiding loops resulting in a backbone graph that is thought reflect the major qualitative properties of connectivity while allowing a better comparison across age groups by avoiding the degree distribution confound. EEG was recorded in a large (N=1500) population-based sample aged 5 to 71 years. Connectivity was assessed using Phase Lag Index to reduce effects of volume conduction. As previously reported, connectivity increased from childhood to adolescence, continuing to grow nonsignificantly into adulthood decreasing only after ~30 years of age. Leaf number, degree, degree correlation, maximum centrality from the MST graph indicated a pattern of increased integration and decreased randomness from childhood into early adulthood. The observed development in network topology suggested that maturation at the neuronal level is aimed to increase connectivity as well as increase integration of the brain network. We confirm that brain network connectivity shows quantitative changes across the life span, and additionally demonstrate parallel qualitative changes in the connectivity pattern.

## Introduction

The brain is a complex network of highly connected brain areas under constant pressure for optimal performance. Describing the brain network using graph theoretical parameters has proven useful providing biomarkers for disease(Heuvel et al., 2010; Menon, 2011; Stam et al., 2009; Tijms et al., 2013; Zhao et al., 2012); In addition, it provides a theoretical underpinning for what might constitute optimal performance in an optimal network organization(Bullmore and Sporns, 2009; de Haan et al., 2009; Sporns, 2014; Stam, 2014a). Ontological development may show similar pressure for increased optimal organization. Anatomically, the human brain shows large changes on a global scale (Casey et al., 2000; Courchesne et al., 2000; Giedd et al., 1999; Lenroot and Giedd, 2006; Paus et al., 2001; Westlye et al., 2010), on the intermediate scale of distinct brain areas (Gogtay et al., 2004; Lenroot and Giedd, 2006; Paus, 2005; Shaw et al., 2006), but also on the neuronal micro scale (Huttenlocher, 1979; Huttenlocher and Dabholkar, 1997; Huttenlocher and de Courten, 1987). These anatomical changes are accompanied by changes on a functional level, as measured using fMRI and M/EEG. The resting state networks seem largely in place by the age of two (Fransson et al., 2010) but also show clear development by increasing (long-distance) connectivity as evidenced both from fMRI (Fair et al., 2009, 2008; Power et al., 2010) and M/EEG studies (Courchesne et al., 2000; Hanlon et al., 1999; Smit et al., 2012). Comparing young to older adults, modularity decreases for longer connectivity distances and across networks (Meunier et al., 2009), and in aging, connectivity decreases in strength (Smit et al., 2012).

Functional methods of determining connectivity use either direct (EEG, MEG) or indirect (fMRI BOLD) measures of correlated neuronal activity to derive coupling strength between brain areas. The high temporal resolution of M/EEG may be particularly useful for estimating short duration networks that appear and disappear on second scale (“fragile binding”). On a larger temporal scale, fMRI can also detect connectivity, as illustrated most clearly by the resting state networks (Damoiseaux et al., 2006). Recent work has shown that both the fMRI and M/EEG based resting state activity share a common ground (Britz et al., 2010; Mantini et al., 2007; Musso et al., 2010). Our previous investigations showed that connectivity showed substantial change over time closely following anatomical developmental curves of white matter (Boersma et al., 2010; Smit et al., 2012, 2010). Moreover, connectivity correlated with white matter volume (Smit et al., 2012). When connectivity matrices from EEG were converted to graphs and analyzed following Watts and Strogatz (Watts and Strogatz, 1998) global network efficiency showed similar correlations with white matter and protracted development from childhood into young adulthood (Smit et al., 2012).

MEG and EEG recordings are subject to volume conduction effects that blur the recorded signals at the scalp or sensor level. Volume conduction is particularly problematic for determining functional connectivity between signals for algorithms like Coherence and Synchronization Likelihood (Nunez et al., 1997; Nunez and Srinivasan, 2006). For this reason, Stam et al. (Stam et al., 2007) proposed the Phase Lag Index (PLI), which reduces the effect of volume conduction by ignoring zero and 180* phase differences between pairs of signals. The PLI algorithm inspects the instantaneous phase of a signals oscillation (e.g., alpha oscillations) by computing the Hilbert transform, instantaneous phase being then determined as the angle of the complex valued signal:

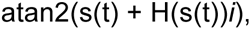

where s(t) is the signal over time t, and H the Hilbert transform, i is sqrt(-1), and atan2 is the arctangent considering the sign of the real and imaginary inputs to return positive or negative angles. Next, the phase difference is computed between pairs of signals as the difference in instantaneous phase and restricted to the [-pi, pi] range. If the distribution of phase differences is symmetric around zero, this may be evidence for spurious connectivity due to volume conduction. Deviances from a symmetric distribution must be due to dependency between sources (direct or indirect). Flat distributions show no evidence for connectivity, spurious or not spurious. Our first aim is to establish whether average connectivity, as well as the graphs derived from connectivity matrices, still show the strong developmental effects that we have reported earlier (Boersma et al., 2013, 2010; Smit et al., 2012, 2010).

A second limitation of previous studies may be that the comparison of networks across the different age groups is problematic, as networks have different average connectivity and degree(van Wijk et al., 2010). Although the use of graph parameters compared to those of randomized graphs is often thought to remove much of these comparability problems, this may only partly be the case(van Wijk et al., 2010). The use of the Minimum Spanning Tree (MST) graph might provide additional information over the use of thresholded or weighted graphs (Boersma et al., 2013; Stam, 2014b; Stam et al., 2014). MST graphs are connected graphs constructed from weighted, undirected connectivity matrices in such a way that they i) are fully connected, and ii) do not form loops, thus forming a “backbone” tree-like graph. Table 1 describes the MST graph parameters that we derived from the MST graphs. Figure 1 shows how graph characteristics can be scaled as a function of the leaf number. It has been argued that optimal network function is a trade-off between small diameter (i.e., the MST network reveals that the underlying brain network is compact) but not dependent on a single hub-node (which has very small diameter but may be less resilient (Albert et al., 2000; Stam, 2014b; Stam et al., 2014). Increased integration of the network may manifests itself as a move from low to high leaf number in MST graphs. How MST graphs relate to optimization of the brain is a matter of ongoing investigation (Stam, 2014b; Stam et al., 2014). However, it has been shown that MST parameters are altered in Parkinson’s disease, epilepsy, Alzheimer’s disease, and near brain tumors (Dubbelink et al., 2014; Tewarie et al., 2014; van Dellen et al., 2014); for an overview, see (Stam, 2014a).

**Table 1.** MST graph parameters and their description

| MST graph parameter | Abbreviation. | Description |
| --- | --- | --- |
| Leaf number | LN | Number of end nodes (i.e., nodes with degree $k=1$) represents the dimension from linear to star graph (figure 1) |
| Diameter | DI | Largest in the set of shortest paths between all possible pairs of nodes. |
| Betweenness Centrality | $BC_{max}$ | Maximum value of the number of shortest paths passing through the nodes |
| Eigenvector centrality | $EC_{max}$ | Maximum value of the loadings on the first principal component of the graph. |
| Maximum degree | $K_{max}$ | Largest degree in the graph |
| Tree hierarchy | TH | Tradeoff between diameter and maximum degree |
| Degree Correlation | DC | Correlation between the degrees of pairs of connected nodes |

**Figure 1.**
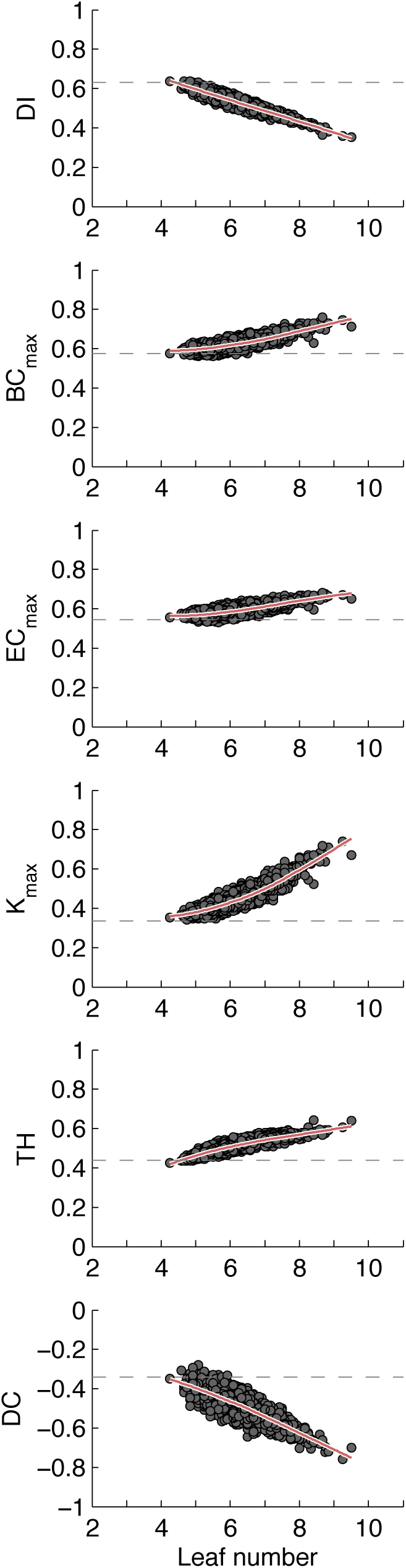
MST graph parameters covary with leaf number (LN). From left to right on the x-axis increased LN indicate increased hierarchical order and integration in the network. LN ranges from 2 (a linear configuration) to 11 (a star-like configuration) for a 12-vertex network, expressed here as a proportion from 0 to 1. Each plot contains the average MST graph parameter values plotted against average LN from the actual PLI based EEG networks. The dashed line Average diameter decreases linearly as a function of LN, as does degree correlation. The centrality-type measures (BC_max_, EC_max_, and K_max_) all show increases with LN with upward curve. Designed as a tradeoff measure, TH increases but with a downward slope for higher leaf numbers, thus penalizing the star-like configurations with extreme K_max_ values. Almost all graphs showed a more star-like organization than the average of random graphs.

In sum, we investigated whether the observed increase in network order remains after connectivity has been established with a measure less sensitive to volume conduction. Next, we investigated whether the MST graphs from connectivity networks provide a similar picture of increased order with maturation. Finally, while our previous analyses focused on a narrow age range in childhood (5 and 7 year olds (Boersma et al., 2013)), here we provide data on subjects aged 5 – 71 from multiple large longitudinal EEG datasets covering adolescence and adulthood. Using these data we will establish whether the previously reported increase in MST order in childhood continues up to adolescence and is maintained during adulthood.

## Methods

### Subjects and procedure

Data were collected as part of a study into the genetics of brain development and cognition. A total number of 1675 individuals (twins and additional siblings) accepted an invitation for extensive EEG measurement. For the present analyses, EEG data recorded during 3–4 minutes of eyes-closed rest were available from six measurement waves with ages centered approximately around 5, 7, 16, 18, 25, and 50 years. Part of these consisted of longitudinal measurements at two ages (5–7 and 16–18 years). In addition, some of the subjects aged 16–18 years were invited back for measurements at age 25. In total, this study incorporated 2540 EEG recordings. After data cleaning, 2137 recordings were available. The structure of the final subject set after data cleaning used in the present study was 331, 368, 418, 380, 350, and 290 respectively for the six measurement waves, which included 294 longitudinal observations between 5 and 7, 374 between 16 and 18, 96 between 18 and 25, of which 95 with measurements at three waves 16, 18, and 25.

Ethical permission was obtained via the “subcommissie voor de ethiek van het mensgebonden onderzoek” of the Academisch Ziekenhuis VU (currently METc of the VU University Medical Centre). All subjects (and parents/guardians for subjects under 18) were informed about the nature of the research. All subjects or parents/guardians were invited by letter to participate, and agreement to participate was obtained in writing. All subjects were treated in accordance with the Declaration of Helsinki.

### EEG acquisition

The childhood and adolescent EEG were recorded with tin electrodes in an ElectroCap connected to a Nihon Kohden PV-441A polygraph with time constant 5 s (corresponding to a 0.03 Hz high-pass filter) and lowpass of 35 Hz, digitized at 250 Hz using an in-house built 12-bit A/D converter board and stored for offline analysis. Leads were Fp1, Fp2, F7, F3, F4, F8, C3, C4, T5, P3, P4, T6, O1, O2, and bipolar horizontal and vertical EOG derivations. Electrode impedances were kept below 5 k**Ω**. Following the recommendation by Pivik et al. (Pivik et al., 1993), tin earlobe electrodes (A1, A2) were fed to separate high-impedance amplifiers, after which the electrically linked output signals served as reference to the EEG signals. Sine waves of 100 μV were used for calibration of the amplification/AD conversion before measurement of each subject.

Young adult and middle-aged EEG was recorded with Ag/AgCl electrodes mounted in an ElectroCap and registered using an AD amplifier developed by Twente Medical Systems (TMS; Enschede, The Netherlands) for 657 subjects and NeuroScan SynAmps 5083 amplifier for 103 subjects. Standard 10-20 positions were F7, F3, F1, Fz, F2, F4, F8, T7, C3, Cz, C4, T8, P7, P3, Pz, P4, P8, O1 and O2. For subjects measured with NeuroScan Fp1, Fp2, and Oz were also recorded. The vertical electro-oculogram (EOG) was recorded bipolarly between two Ag/AgCl electrodes, affixed one cm below the right eye and one cm above the eyebrow of the right eye. The horizontal EOG was recorded bipolarly between two Ag/AgCl electrodes affixed one cm left from the left eye and one cm right from the right eye. An Ag/AgCl electrode placed on the forehead was used as a ground electrode. Impedances of all EEG electrodes were kept below 5 k**Ω**, and impedances of the EOG electrodes were kept below 10 k**Ω**. The EEG was amplified, digitized at 250 Hz and stored for offline processing.

### EEG preprocessing

We selected 12 EEG signals (F7, F3, F4, F8, C3, C4, T5, P3, P4, T6, O1, O2 and both EOG channels) for further analysis as the set with the most complete match between the different measurement waves/cohorts.

All signals were broadband filtered from 1 to 35 Hz with a zero-phase FIR filter with 6dB roll-off. Next, we visually inspected the traces and removed bad signals. Note that for the network analysis a full set of EEG signals was required and therefore any rejected EEG channel resulted in the loss of that subject. Next, we used the extended ICA decomposition implemented in EEGLAB to remove artifacts, including eye movements, and blinks. After exclusion of components reflecting artifacts, the EEG signals were filtered into the alpha (6.0 to 13.0 Hz) frequency band. The peak alpha frequency developed from 8.1 Hz at age 5 to 9.9 Hz at age 18, after which a slow decline to 9.4 Hz was observed at around 50 years. The lower edge of the alpha filter was set such that alpha oscillation of all subjects was included from ~2.0 Hz below the lowest peak frequency to ~3.0 Hz above the highest average peak frequency.

### Connectivity

Connectivity was calculated using the Phase Lag Index (PLI). For a detailed description we refer the reader to Stam et al., (Stam et al., 2007). In short, the phase lag index inspects the distribution of phase differences between pairs of signals ***X****={X_1_,X_2_,X_3_…X_N_}*. First, signals in ***X*** are filtered for oscillations in the frequency band of interest. Next, instantaneous phase for a signal *X*_*i*_ is established using the Hilbert transform *H* (*X_i_*)(*t*)), as described above. Phase difference between signals *n* and *m* is then *Δϕ*(*t*) *= ϕ_n_* (*t*) − *ϕ_m_*(*t*) for *n* ≠ *m*. Next PLI is calculated as

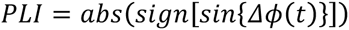

for *Δϕ* modulated within the range −*π* and *π*.

### Graph analysis

MST Graphs were created with the Kruskal algorithm applied to the PLI connectivity matrices. Next, we derived parameters described in table 1 from these graphs using a variety of MATLAB algorithms including standard MATLAB code, the MIT graph toolbox (http://strategic.mit.edu/downloads.php?page=matlab_networks), the brain connectivity toolbox (Rubinov and Sporns, 2010), and custom scripts. We performed the same analysis on 1000 random graphs by creating symmetric matrices with random numbers on a (0, 1) interval. We extracted the same graph parameters (Table 1) and averaged these across the 1000 graphs.

### Statistics

The effect of age was determined in several ways. First, we created developmental plots (scatterplots) from connectivity and each of the MST graph parameters on age. Next, local nonlinear weighted regression trends were fitted (loess, on 65% window size second order polynomials). 95% confidence intervals were obtained using a bootstrap with 10000 repeats. Since some observations are nested within family, the bootstrap was based on the independent unit (family) rather than individual. Note, however, that the bootstrap was not used to establish significance. To test significance of developmental trends, we estimated different fixed effect models. First, linear, quadratic, and cubic trends were fitted to the dataset, which we tested for significance. Because the complex structure of the data including repeated measures and family dependencies, which even extended across the different age groups (siblings of twins might fall into a different age category than the proband twins), we used Generalized Estimating Equations (GEE) to obtain p-values. GEE with the exchangeable correlation matrix estimates a single correlation across residuals within clusters (i.e., family number). Even though the residual matrix is in fact more complex than the single estimated working correlation (for example, within-subject correlations and MZ twin correlations are expected to be higher than other within-family correlations), the robust SEs are not affected by this misspecification (Minică et al., 2014).

Second, we defined nine age-groups using the following boundaries specified in years: 4.9 – 6.0, 6.0 – 7.4, 13.0 – 16.6, 16.6 – 20.0, 20.0 – 25.0, 25.0 – 35.0, 35.0 – 45.0, 45.0 – 57.5, and 57.5 and older. These were tested in a pair-wise fashion for significance with FDR correction for the n(n-1)/2 comparisons (36 at n=9) tested at q=0.025 (Benjamini and Hochberg, 1995). This level of q was chosen to accommodate the dimensionality of the data, which showed a clear two-dimensional structure (see results).

## Results

### Increased order with increased leaf number

Figure 1 shows the dependency of MST graph parameters on LN. Each point in the scatterplot represents the MST graph values of a single individual. Note that most values fall close to the polynomial regression. A second order polynomial fit was significant in all cases. The centrality measures and K_max_ showed a positive relation with upward curve as expected from random graph simulations (Boersma et al., 2013). TH also showed a positive dependence on LN, but with a downward curve, thus setting a limit to the effect of LN on TH. DI decreased with increasing leaf number in a linear fashion, which may be expected since maximum and minimum values for DI may be derived analytically from LN (Stam, 2014b). DI will lie between *DI*_max_ = *N* − *L* + 1 and *DI*_*min*_ = *2* (*N* − 1)/*L*.

### PLI connectivity shows an inverted-U development

Figure 2 (top row) shows the results of average connectivity developing over age. PLI connectivity showed a pattern of development similar to those reported previously based on a different measure of connectivity (Smit et al., 2012). Left column (A) shows the development with loess fit (50% window size, 2^nd^ order fit). The data points reveal large individual differences. The middle column (B) reveals changes from childhood to early adulthood in average PLI, a decrease from 16 to 25 after which a plateau was reached. Bootstrapping confidence intervals suggest that significant changes are present in the data, especially from childhood to adolescence. To test significance we compared age groups in pairwise manner. Significant increases were found from childhood to adolescence, but also between ages 5 and 7. A significant decline in connectivity was observed in the 50+ age group compared to adolescent and other adult age groups (except age group ~22).

**Figure 2.**
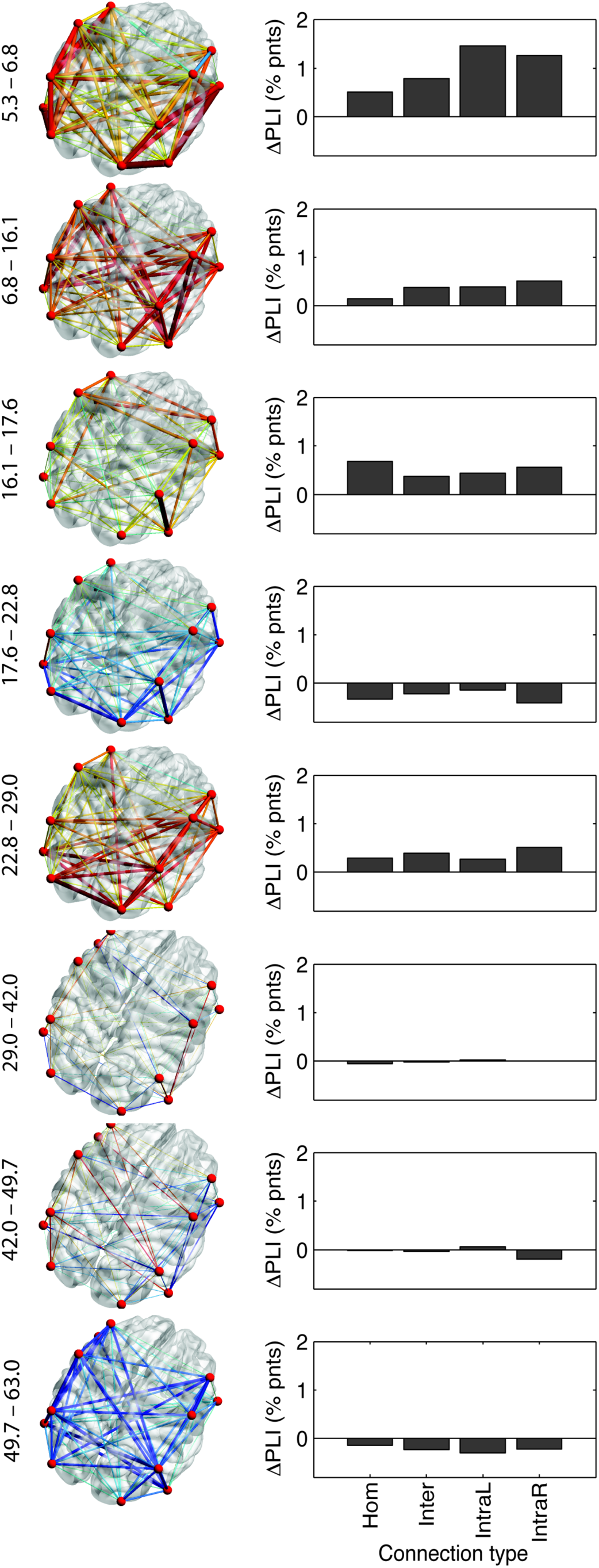
Localized development of connectivity strength. (Left) 3D heaplots of average change in PLI per year from one age group to the next (age groups 5, 7, 16, 18, 22, ~30, ~40, ~50, 57.5+) from an elevated right posterior viewpoint. The location of maximal development is not stable, but changes with age. (Right) Separating edges into intra-hemispheric left and right (IntraL, IntraR), contralateral homologues (Hom) and other cross-hemisphere (Cross) connections showed that childhood was marked by a clear Intrahemispheric increase of connectivity, while later age groups showed no such strong prevalence, or stronger increases in contralateral homologues (within adolescence).

Our previous report on the same sample used Synchronization Likelihood (SL) as a measure of functional brain connectivity. The current results show remarkable similarities. Both PLI and SL showed strong development from childhood to adolescence with effect sizes over r>0.40 comparing age group 5 with other ages. PLI showed peak value at age 40 (Figure 2C), where the SL peaked at around age 50. This suggests that the previous results were quite robust against effects of volume conduction and common reference. However, the current results also differed from those reported previously. Connectivity measured using SL showed a continuous increase up to the peak age, whereas PLI connectivity showed a decrease from age 18 to ~22.

### Connectivity patterns change with age

Although average PLI connectivity may change similarly across different age groups, localized differences may still occur, leading to different network types. We assessed the connectivity between all possible pairs of signals and calculated change across the 8 age groups in rate per annum. 3D headplots were constructed using BrainNetViewer with approximate locations of the electrodes (see Figure 2A). The thickness of the edges were rescaled to average increase per annum making it comparable across headplots. Red colors indicate increase, blue decrease, and green indicates no change.

Changes within childhood were largely limited to intrahemispheric connections (figure 2B). Homologous (left-right lateralized) electrode positions and other interhemispheric connections showed low PLI connectivity increase. From childhood to adolescence both inter- and intrahemispheric connections showed increases, but homologues still showed less PLI change. In adolescence, a change is seen with homologues reaching the largest change. In later ages, reduction in connectivity strength is clearest in interhemispheric connections (other than homologues). In sum, the changes during childhood, adolescence and middle-age show remarkable differences in topology. Clearly, the brain does not simply change connectivity but changes the overall pattern of connectivity.

### An increasingly integrated network

Figure 2 shows the development of MST graph parameters as scatterplot with loess fit (A), bootstrap of the loess fit with 95% confidence intervals (B), and pairwise testing of significance across age groups (C). Network parameters showed developmental trends highly comparable to connectivity. Cubic curves were not significant (absolute robust z<1.4, *ns*). All quadratic terms were significant (absolute robust z>6.44, p<1.2E-10) with all parameters showing inverted-U shapes—except MST diameter showed a U curve as expected.

The brain network of children showed lower leaf number, indicating a more line-like / less integrated organization. Increasing age resulted in increased leaf number and a correspondingly increased BC_max_, EC_max_, K_max_. TH changed similarly in an inverted-u shape. DI and DC decreased. These findings are consistent with an increasingly star-like organization and increased integration. The comparison of most measures showed significant change from 5 years of age to adolescence/adulthood with highly significant values (p<0.0001, and p<0.001 compared to age ~40 for DC and EC_max_). Age 7 showed a similar pattern. Both ages 5 and 7 generally showed no significant difference with the oldest age group (>57.5).

Older age (57.5+) was marked by significant decrease for many parameters, although the effects were not very strong (p<0.01). LN and TH decreased with older age compared to ages ~30 to ~50. Connectivity decreased only compared to age ~40 (p<0.01). The centrality measures showed less consistent decrease, possibly due to a noisier variation.

### Principal components reveal partly separate sources of variation

Since the developmental trends of connectivity and MST graph parameters showed markedly similar paths, we subjected the correlation matrix of four different measures for connectivity (homologous contralateral connectivity, other interhemispheric connectivity, intrahemispheric connectivity left and right) and six MST based graph parameters (LN, DI, BC_max_, EC_max_, K_max_, DC) to an eigenvalue decomposition after selecting one random subject per family and regressing out the effects of age and sex. TH was excluded since it was based on two other parameters and therefore does not add information to the correlation matrix. Scores for DI and DC were inversed so as to enforce positive correlations. Figure 3 shows the results. The correlation matrix shows a clear clustering of connectivity versus MST graph parameters. The highest eigenvalue of 5.85 explained 58.5% of the variance, the second highest was 2.09 (20.9% variation). Both the correlation matrix (Figure 3A) and the scree plot (Figure 3C) strongly suggest a two-factor solution. After varimax rotation MST parameters loaded strongly on the first component and PLI connectivity measures on the second (Figure 2B). Figure 2D shows that the two components show different developmental patterns, with a much clearer U-curve for MST graph parameters, while PLI connectivity shows a decrease from adolescence to young adulthood. For these reasons, we conclude that MST graphs parameters and PLI based connectivity largely reflect different sources of variation in brain function with different developmental curves.

**Figure 3.**
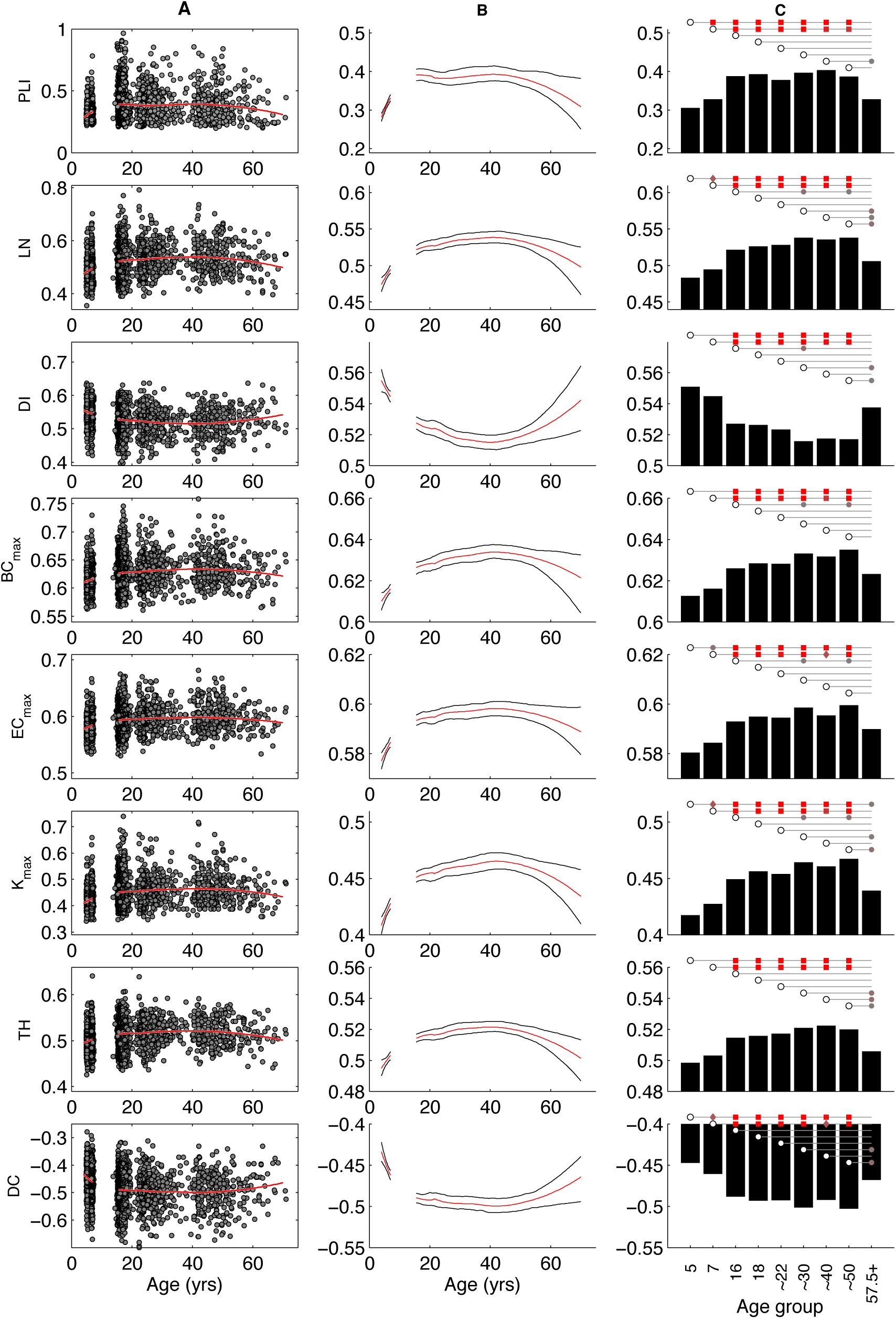
Age development plots for PLI and MST network parameters of EEG alpha oscillations (6.0 – 13.0 Hz). (A) Large individual variation around loess smooth (70% width, 2^nd^ order fit) was observed. Most parameters showed (inverted) U curved development. (B) The same loess smooth was used in a clustered bootstrap function with families as sampling units, thus keeping the residual correlations on average intact. Shown are 95% CI range and median values obtained in the bootstrap. (C) Group-wise comparison was corrected for residual correlation with robust SEs (Generalized Estimating Equations, see methods) and FDR corrected for the n*(n-1)/2 comparisons at n=9 and with q=0.025 to correct for the dimensionality of the data, which was set to 2 as per figure 4. Pairwise comparison was significant when an open circle is connected to a grey/red marker. For example, age group 5 differed significantly in PLI connectivity strength from ages 7, 16, 18, 22, and 30. Squares indicate the strongest effect (corrected-*P*<0.0001) followed by diamonds (corrected-*P* <0.001) and circles (corrected-*P* <0.01). Color saturation indicates –log10(*P*), with grey values for corrected-*P* = 0.05 ranging to bright red for corrected-*P* = 10^-6^.

**Figure 4.**
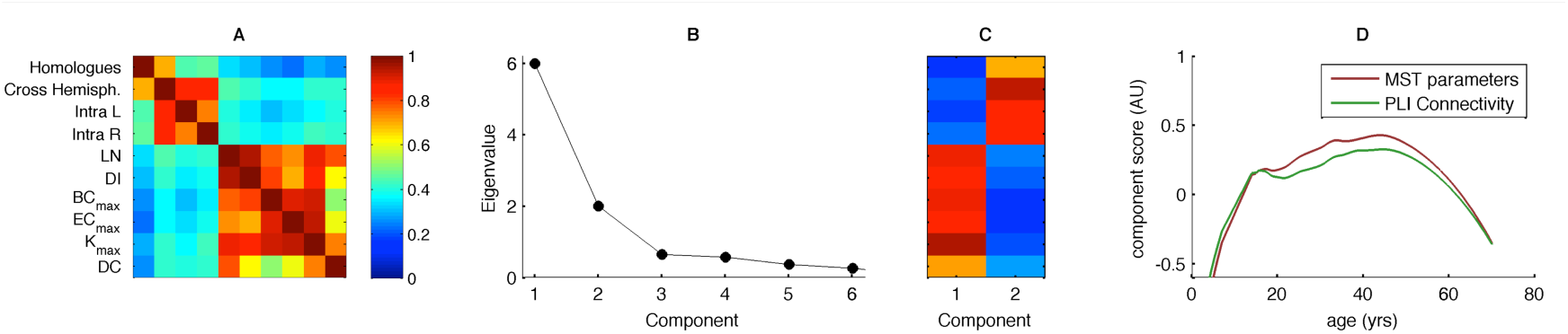
Principal Components Analysis of connectivity scores separated into contralateral homologues (Hom), other cross-hemispheric connections (Cross) intra-hemisphere left (Intra L) and intra-hemisphere right (Intra R), and MST graph measures. Note that TH was excluded as this measure is fully based on two other graph parameters (DI and Kmax). Positive correlations of DI and DC with other parameters was enforced by reversing scores. (A) The correlation matrix (corrected for age and sex) suggested two clusters, one for the four connectivity types, and one for the MST graph parameters. (B) The,scree plot also strongly suggested two separate sources of variation. (C) The loading pattern for varimax-rotated two-component extraction showed clear separation of the connectivity and MST graph measures. (D) The varimax rotated factor scores (corrected for sex) showed different developmental paths, suggesting that development differentially affects connectivity and MST graph parameters.

## Discussion

Our aim was to investigate whether the increased integration of the network observed from 5 to 7 years of age extends into adolescence and adulthood. The large and highly significant differences found in graph parameters and connectivity between childhood and adolescence/adulthood suggest that this is the case. We established that life-span development of average connectivity between pairs of scalp-recorded signals closely mimic those reported previously (Smit et al., 2012). By using the PLI (Stam et al., 2007)—a measure that ignores volume conduction—we have found support that our previous findings using synchronization likelihood (Stam and van Dijk, 2002) have not been spurious. We hypothesize that the sparse electrode layout in our previous report may have been protective against detecting false synchronization (Smit et al., 2012).

Average connectivity measured with PLI showed strong increases within childhood and from childhood to adolescence. Several findings in the extant literature suggest that this increase in EEG functional connectivity depends on maturation of white brain matter, including myelinization. For example, it has been found that interhemispheric EEG connectivity measured by coherence has been related to DTI diffusivity (Teipel et al., 2009), and to T2 relaxation times in white matter in head injury, which arguably is related to neuronal membrane lesion (Thatcher et al., 1998). In addition, we have previously found that developmental curves for connectivity are highly consistent with the protracted development of white matter development: both connectivity and White Matter Volume (WMV) showed peaks in middle age (Allen et al., 2005; Bartzokis et al., 2001; Benes et al., 1994; Good et al., 2002; Walhovd et al., 2005a, 2005b; Westlye et al., 2010). Moreover, a moderate correlation was found between WMV and connectivity. Because PLI reduces the effects of spurious connectivity in the brain based on volume conduction and common reference, these results seems to further strengthen the idea that functional connectivity in the resting-state reflects the strength of anatomical connectivity between distant brain areas.

Arguably, MST graphs are more comparable across groups than thresholded graphs (Stam, 2014b; van Wijk et al., 2010). Graph parameters derived from the MST graph showed evidence for change in the level of integration. All MST parameters show an inverted-U curve (and a U curve for diameter). The backbone graph in human brain activity moved from a line to a more star-like configuration during development. In later age, a return to a more line-like configuration was found. For all but the centrality measures, these resulted in significant drops for age group 57.5+ compared to ages 30 and 50. Importantly, principal components analysis showed that MST graph parameters reflected different sources of variation compared to PLI connectivity. Clearly, not just the average connectivity, but the connectivity *pattern* changes. Note that we observed that MST graph parameters showed a more star-like configuration than random graphs (Figure 1). In this sense, the observed developmental changes showed a move from random networks towards more integrated networks, and more random networks in later life. This, too, is consistent with previous observations of life-span development in the same sample (Boersma et al., 2010; Smit et al., 2012, 2010).

The results make MST graph parameters highly suitable as biomarkers for development in early life and cognitive decline associated with older age. Follow-up studies could target the genetic variants that have been linked to neuronal change such as myelination. Additionally, studies could investigate how genetic variants exert their influence in cognitive decline or Alzheimer’s disease (e.g., APOE, CLU/APOJ, and PICALM (Harold et al., 2009; Hollingworth et al., 2011; Lambert et al., 2013). Carriers of the APOE \epsilon4 allele have an increased risk for forming beta-amyloid plaques; during prion-like aggregation, damage to neurons is done by oxidative stress, resulting in brain atrophy. This loss significantly reduces the number of neurons available for connectivity such as seen in MCI and AD (Jelic et al., 1997; Tóth et al., 2014), but may also result in the loss of integration in the MST network. Likewise, clusterin (CLU/APOJ) is involved in the clearance by binding to beta-amyloid resulting in variability in neurodegeneration (Desikan RS et al., 2014; Mengel-From et al., 2013) and could have similar effects on connectivity and connectivity patterns. PICALM highights the need to investigate inflammatory pathways (Perry et al., 2010). From the current results we expect that connectivity loss will prove to be nonrandom, resulting in reduced integration due to specifc atacks on central nodes (see also (He et al., 2009; Stam et al., 2009)).

In developmental neurobiology, the dichomotmy into long and shorter projection distances may be essential. In an fMRI study, it was shown that decreased short range connectivity concurs with increased long-range connectivity. Local connections in a cognitive control network become less diffuse with development from 10 to 22 years of age, which is accompanied by increased long distance functional connectivity (Kelly et al., 2009). Similar findings of changes in (long-distance) connectivity have been reported (Dosenbach et al., 2010; Fair et al., 2009; Supekar et al., 2009). The present results extend these findings in showing that from childhood to adulthood brain networks move from less to more integrated graphs (figure 2). Since network parameters may be relevant predictors of cognitive performance (Micheloyannis et al., 2006; Tewarie et al., 2014; van den Heuvel et al., 2009) and are disrupted in neurological disorders (Stam et al., 2014, 2009; Tewarie et al., 2014; van Dellen et al., 2014), we can hypothesize that the increasingly integrated network topology is essential to the large developmental changes in human cognitive performance during the same period. Indeed, a more integrated network was predictive of better cognitive performance in MS patients (Tewarie et al., 2014). Cognitive performance correlated with a larger decrease in network integration in Parkinson’s patients (Olde Dubbelink et al., 2014). Whether these findings generalize to the normal population may be addressed in future investigations.

In conclusion, brain connectivity measured by the PLI shows large changes over the lifespan. These changes largely corroborate the earlier findings that connection strength increases during development (Hagmann et al., 2010; Smit et al., 2012, 2010). Since PLI is less sensitive to volume conduction by ignoring the zero and pi phase differences between signal pairs (Stam et al., 2007), developmental changes are therefore unlikely to reflect changes in conductive properties across age groups. The use of the minimum spanning tree backbone graph aimed to solve the problem that graph measures may not be compared across different sizes and degree distributions (van Wijk et al., 2010). However, MST graphs confirmed that brain matures across the lifespan and shows changes in structure both in the development in childhood and during ageing later life. These findings corroborates our earlier findings that the network shows reduced randomness from childhood to young adulthood (Boersma et al., 2013, 2010; Schutte et al., 2013; Smit et al., 2012, 2010).

## Acknowledgements

This work was supported by grants Twin-family database for behavior genetics and genomics studies (NWO 480-04-004) to D.B., Genotype/phenotype database for behavior genetic and genetic epidemiological studies (NWO 911-09-032) to D.B., European Research Council (ERC-230374) to D.B., BBR Foundation (NARSAD) Young Investigator grant 21668 to D.S., NWO/MagW VENI-451-08-026 to D.S., VU University VU-USF 96/22 to D.B., Human Frontiers of Science Program RG0154/1998-B to D.B. and E.d.G., Netherlands Organization for Scientific Research, NWO/SPI 56-464-14192 to D.B.

